# Adaptive immunity shapes innate and epithelial cell landscapes by silencing tonic IFN-γ in innate lymphoid cells during homeostasis

**DOI:** 10.1101/2024.09.19.613869

**Authors:** Ousséma Mejri, Li Ma, Amneh Aoudi, Ossama Labiad, Stanislas Mondot, Ramdane Igalouzene, Clément Codan, Hector Hernandez-Vargas, Noémi Rousseaux, Nicolas Lapaque, Stéphane Paul, Thierry Walzer, Julien C. Marie, Saïdi Soudja

**Affiliations:** Centre International de Recherche en Infectiologie, INSERM, U1111, CNRS, UMR5308, Ecole Normale Supérieure de Lyon, (ENS Lyon), Université Claude Bernard Lyon 1 (UCBL), Lyon, France; Centre de Recherche en Cancérologie de Lyon, INSERM U1052 CNRS UMR5286, Centre Léon Bérard, Université Claude Bernard Lyon 1 (UCBL), Lyon, France; Université Jean Monnet, Saint-Etienne, France; Université Paris-Saclay, INRAE, AgroParisTech, Micalis Institute, Jouy-en-Josas, France; Genomics Consulting, 69500 Bron, France

## Abstract

During homeostasis, innate and epithelial cells undergo continuous maturation, shaped by microbial, molecular and cellular interactions. While these cells influence adaptive immunity during steady-state conditions, the reciprocal homeostatic role of adaptive immune cells in the maturation and function of innate and epithelial cells remains underexplored. Here, through computational approaches and murine models, we establish that adaptive immunity shapes innate immune and epithelial homeostatic landscapes. Mechanistically, adaptive immunity acts as a brake on type 1 polarization and diversification of innate lymphocytes (ILCs). Without adaptive immunity, the innate cellular interactions within the mesenteric lymph nodes are dominated by an interferon-gamma (IFNγ) signaling network. Moreover, the innate immune and epithelial cells follow distinct phenotypic and functional trajectories, including ILCs acquiring myeloid markers, monocytes adopting an inflammatory path, and colonic epithelial cells expressing altered antimicrobial peptides and losing Paneth-like features. Depleting ILCs, abolishing IFNγ signaling, or restoring adaptive immunity reduces these IFNγ-driven changes. Thus, our findings highlight a homeostatic function of adaptive immunity in modulating innate and epithelial cell communication, diversity, composition, function, and differentiation, notably by limiting ILC-derived IFNγ.

## Introduction

Epithelial cells and innate immune cells, including monocytes (MOs), dendritic cells (DCs), macrophages (MACs), innate lymphoid cells (ILCs), and NK cells, constitute the first line of defense and play a key role in the initial detection and protection against pathogens. However, innate immune cells have to undergo a lengthy maturation and preparation during homeostasis to be fully operational (Bosteels et al., 2023; Dalod et al., 2014; Lutz and Schuler, 2002). This “homeostatic shaping/maturation” process requires microbial, metabolic, molecular, and cellular interactions.

Numerous studies have demonstrated how commensal microorganisms, such as segmented filamentous bacteria (SFB) and *Akkermansia muciniphila*, influence the development and function of immune and epithelial cells (Ivanov et al., 2008; Rie Gaboriau-Routhiau et al.; Ansaldo et al., 2019). During homeostasis, the microbiota provides basal and tonic signals that continuously shape immune and epithelial cells. These instructive signals are transmitted through metabolites, toll-like receptor ligands, and microbiota-derived DNA and are translated into a tonic interferon (IFN) signal (Pott and Stockinger, 2017; Abt et al., 2012; Van Winkle et al., 2022). IFNs exist in three types: I, II, and III. While type I and III IFNs have well-established roles in microbiota-driven homeostatic immune and epithelial cell shaping, type II IFN (IFNγ) has not typically been associated with this process. The tonic homeostatic IFN-signaling, mainly attributed to type I and III, pre-arms innate immune cells and epithelial cells with IFN-stimulated genes (ISGs), many of which encode anti-microbial molecules. Disrupting this tonic IFN signal compromises the host defense against pathogens (Pott and Hornef, 2012; Pott et al., 2011; Saskia Erttmann et al., 2022; Schaupp et al., 2020). Therefore, the microbiota-mediated homeostatic shaping of innate immune and epithelial cells via IFNs is key for protecting the organism.

Other parameters are involved in “homeostatic shaping”, such as cellular interactions. Upon continuous stimulation by the microbiota, myeloid cells produce a tonic type 1 IFN, which primes innate immune and adaptive immune cells during homeostasis (Pott and Hornef, 2012; Pott et al., 2011; Saskia Erttmann et al., 2022; Schaupp et al., 2020; Ganal et al., 2012). Additionally, epithelial cells from the small intestine, upon attachment of SFB, secrete serum amyloid A (SAA), which directs DCs to differentiate naïve CD4 T cells into IL17-producing cells (Ivanov et al., 2008; Rie Gaboriau-Routhiau et al.; Brabec et al., 2023). These examples illustrate the homeostatic role of innate immune and epithelial cells in shaping adaptive immune cell differentiation and function during steady-state conditions. However, whether adaptive immune cells can reciprocally influence innate and epithelial cells under homeostatic conditions remains an open question. Here, we used diverse murine models, computational methods, and single-cell RNA sequencing (scRNA-seq) to determine the impact of the adaptive immune system on the composition, communication, differentiation, and function of innate immune and epithelial cells during homeostasis.

## Results

### 1. Distinct functional and maturation states of myeloid cells without adaptive immunity

To investigate the influence of adaptive immune cells on myeloid cells, we used a reductionist approach comparing myeloid cells isolated from the mesenteric lymph nodes (MLN) of RAG2 knockout (RAG2KO) mice lacking adaptive immune cells to those from wild-type (WT) mice. We employed scRNA-Seq to identify and unbiasedly characterize the cells. Through unsupervised clustering, we identified ten distinct myeloid cell clusters in the MLN. These clusters included six distinct dendritic cell (DC) subsets: a conventional DC1 subset (marked by *Xcr1* and *Clec9a*), two DC2 subsets (marked by *Cd7* and *Cd209*), and three migratory DC subsets (characterized by *Ccr7*, *Fscn1*, *Cd63*, *Nudt17*, and *Ccl22*). Additionally, we identified a MAC cluster (*C1qa*, *Mafb*), a Ly6c+ MO cluster (*Ccr2*, *Fcgr1*), a Ly6c-MO cluster (*Ear2*, *Ly6i*), and a neutrophil cluster (*Cxcr2*, *S100a8*, *S100a9*) (**Fig. 1 A** and **B**). Remarkably, RAG2KO mice showed an increase in Ly6c+ MOs and a decrease in Ly6c-MOs compared to WT mice. Additionally, a specific subset of the DC2 population, termed cDC2a (expressing *Mreg* and *Relb*), was rare in RAG2KO mice, while the MAC population was enriched (**Fig. 1 C**). This result indicates a change in the composition of the myeloid compartment at steady-state conditions.

**Figure 1:**
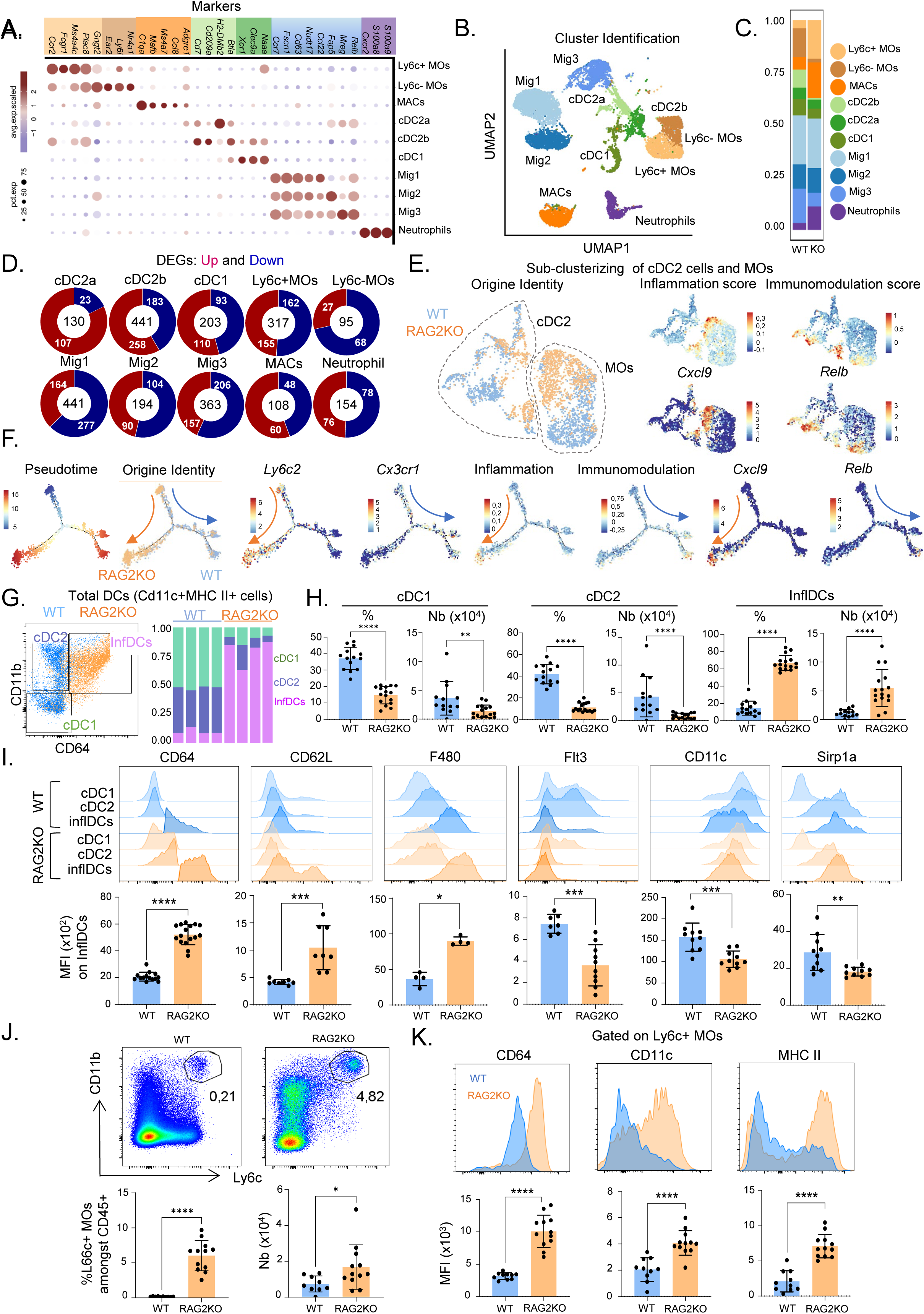
Distinct functional and maturation states of myeloid cells in the absence of adaptive IS. scRNA-seq on sorted myeloid cells of MLN from WT and RAG2KO mice. (A) Dot plots showing the expression of marker genes used to identify myeloid cell subsets. (B) UMAP plot displaying the identified myeloid cell subsets. (C) Bar plot depicting the distribution of each myeloid cell subset based on genotype (WT vs. RAG2KO). (D) Donut plots illustrating the DEGs between WT and RAG2KO mice across various cell types, with the total number of DEGs displayed in the center of each plot (Red DEGs up and blue down in RAG2KO mice). (E) UMAP plots showing sub-clustering by origin (blue = WT, orange = RAG2KO), inflammation score, immunomodulatory score, and the expression of Cxcl9 and Relb. (F) Trajectory inference (Monocle 2) of MO plotted by pseudotime, origin identity, *Ly6c2* and *Cx3cr1* expression, as well as inflammation and immunomodulation scores, and *Cxcl9* and *Relb* expression. (G) FACS dot plot gated on all dendritic cells (CD11c+MHC II+) from WT and RAG2KO mice (WT = blue, RAG2KO = orange), accompanied by a bar plot showing the proportions of cDC1, cDC2, and inflammatory DCs (InflDCs) within the total DC population. (H) Percentage and absolute numbers of cDC1, cDC2 and InflDCs in MLN. (I) FACS histograms of indicated marker expression on cDC1, cDC2 and InflDCs in MLN with mean fluorescence intensity (MFI) values for each marker displayed below (blue=WT and orange= RAG2KO). (J) FACS plot gated on CD45+ cells showing Ly6c+ MOs (CD11b+Ly6c+) in the MLN, along with the percentage and absolute numbers shown below. (K) FACS histograms of CD64, CD11c, and MHC II expression on Ly6c+ MOs, with the MFI values shown below. Data (G-K) represent at least 2 independent experiments and are presented as mean ± SD. Each symbol represents an individual mouse. Data were analyzed by Mann–Whitney tests (\**P* < 0.05, \*\**P* < 0.01, \*\*\**P* < 0.001, \*\*\*\**P* < 0.0001. ns, not significant (*P* > 0.05).

Differential gene expression analysis revealed transcriptional alterations between mouse strains across all myeloid cell populations. Specifically, Ly6c+ MOs, cDC2b, cDC1, and MACs exhibited substantial transcriptional differences at steady-state conditions, with 317, 441, 203, and 108 differentially expressed genes (DEGs) respectively (**Fig. 1 D**). Thus, these findings demonstrate that the absence of adaptive immune cells during homeostasis affects not only the composition of the myeloid compartment but also their transcriptional landscapes.

Given the difficulty in distinguishing between MO and DC2 populations, particularly when DC2 cells are activated (Bosteels et al., 2020), we refined our analysis by sub-clustered DC2 cells and all MOs together. Remarkably, MO and DC2 populations from RAG2KO and WT mice segregated distinctly (**Fig. 1 E**), indicating a large transcriptional divergence. Using an inflammatory score derived from a gene set in the Molecular Signatures Database (MSigDB), we identified clusters with a pronounced inflammatory phenotype in the DC2 and MO compartments. These inflamed clusters were predominantly observed in RAG2KO mice (**Fig. 1E**). A trajectory analysis using Monocle 2 (Trapnell et al., 2014) and TFvelo (Li et al., 2024), conjointly demonstrated that Ly6c+ MOs in RAG2KO mice predominantly differentiate into inflammatory cells expressing *Ly6c2* and the inflammatory chemokine *Cxcl9*. Conversely, in WT mice, Ly6c+ MOs primarily differentiated into Cx3cr1+ Ly6c-MOs with an immunomodulatory profile characterized by the expression of genes such as *Relb* associated with the tolerogenic function of myeloid cells (Wu et al., 2016) (**Fig. 1F** and **S1, A** and **B**). Therefore, our analysis employing different algorithms showed that Ly6c+ MOs in RAG2KO mice adopt a distinct trajectory and an inflammatory state compared to WT mice.

After using flow cytometry (FACS) as a complementary approach, we noticed a significant accumulation of a subset of DCs (CD11c+ MHC II+) expressing the monocytic marker CD64 in RAG2KO mice (**Fig.1 G** and **H**). This subset was termed homeostatic inflammatory DCs (InflDCs) and may include MO-derived DCs and activated DC2 populations (Bosteels et al., 2020). To further characterize the homeostatic inflDCs population, we examined the expression of additional markers. The high expression of CD64, CD62L, and F480, combined with the low expression of typical DC markers such as Flt3, CD11c, and Sirp1a, suggests that the homeostatic inflDCs in RAG2KO mice mainly originate from Ly6c+ MOs (**Fig. 1 I**). Accordingly, the Ly6c+ MOs in RAG2KO mice were abundant and upregulated CD64, CD11c, and MHC II (**Fig.1 J-K**). Therefore, in the absence of adaptive immunity during steady-state conditions, Ly6c+ MOs preferentially differentiate into homeostatic inflDCs.

Given the proposed intrinsic role of Rag genes in NK cells (Karo et al., 2014), we conducted bone marrow reconstitution experiments to delineate whether intrinsic or extrinsic factors drive the functional state alterations in the myeloid compartment of RAG2KO mice. RAG2KO mice were irradiated and reconstituted with a mixture of WT (CD45.1) and RAG2KO (CD45.2) bone marrow. Closer examination, post-reconstitution, showed that both WT and RAG2KO Ly6c+ MOs displayed comparable functional differentiation and distribution patterns. The expression of CD64, CD11c, and MHC II are similar between WT and RAG2KO cells (**Fig.S1 C**). In addition, a similar proportion of homeostatic inflDCs was observed between mouse strains (**Fig.S1 D**). These findings established that myeloid cells from RAG2KO and WT mice undergo similar differentiation paths when exposed to the same environmental conditions. Consequently, without adaptive immunity, extrinsic factors drive myeloid cells to adopt distinct functional and phenotypic states.

### 2. Myeloid cells exhibit a marked interferon signature without adaptive immunity

Gene ontology (GO) analysis showed an enrichment of pathways linked to viral and bacterial responses, as well as type 1 and 2 IFN signaling throughout most myeloid subsets (**Fig. S1, E**). Genes associated with anti-microbial activity (Gbps), class II MHC molecules (*H2-Ab1*, *Ciita*), and immunoproteasome components (*Psmb8*, *Psmb9*) as well as IFN signaling (*Stat1*, *Socs1* and *Irf1*) were upregulated in RAG2KO mice across several myeloid subsets, particularly in Ly6c+ MOs (**Fig. S1, F-G**). Then, we performed an Immune Response Enrichment Analysis (IREA) based on the Immune Dictionary, which delineates immune cell-specific transcriptional responses to distinct cytokines in vivo (Cui et al., 2024). This analysis identified cell-specific responses to type I and type II IFN (**Fig. S1, H**). Thus, without adaptive immunity, myeloid cells display a strong IFNγ signature.

To get an insight into the external factor eliciting this profound myeloid phenotypic and functional alteration in RAG2KO mice, we performed bulk RNA sequencing (RNA-seq) analysis on MLN tissues. We observed an enrichment of inflammatory and type 1 and type 2 IFN signatures in the MLN environment of RAG2KO compared to WT mice (**Fig. S1, I-J**). To formally validate this IFN-biased environment in RAG2KO mice, we measured the level of various cytokines in MLN. While IFNβ were not elevated in MLN of RAG2KO mice compared to MLN of control mice, IFNγ and cytokines known to drive IFNγ production, such as IL12 and IL18 (Okamura et al., 1998), were significantly increased at both transcript and protein levels (**Fig. S1, 1K** and **L**). Consequently, the strong IFN signature exhibited by the myeloid cell population is largely attributed to a tonic IFNγ production within the MLN of RAG2KO mice during homeostasis.

### 3. Diversification and type 1 polarization of innate lymphocytes during homeostasis of innate lymphocytes without adaptive immunity

Next, we analyzed the impact of adaptive immunity on the transcriptome of innate lymphocytes. For this purpose, we performed a scRNA-seq on sorted ILC/NK cells from the MLN (CD45+, CD90+, CD3- and CD19/B220-). Unsupervised clustering revealed 17 distinct clusters after the exclusion of T and B cell contaminants (**Fig. 2 A**). NK cells were distinguished by the expression of *Eomes* and the absence of *Il7r*, whereas the expression of *Il7r* defined other ILCs. Cluster 3 was characterized as ILC2 based on *Gata3* and *Il4* expression, while the presence of *Tbx21* and the absence of *Rorc* identified ILC1. Cluster 13 corresponded to bona fide ILC3, exhibiting high *Rorc*, *Ccr6*, and *Cd4* expression. Clusters 5 and 4 were designated as ILC3/ILC1 subsets due to the co-expression of both ILC3 (*Rorc*, *Kit*, *Ahr*) and ILC1 (*Tbx21*, *Ifng*) markers (**Fig. 2 B** and **C** and **Fig. S2, A**). Unexpectedly, the ILC3 (*Rorc*, *Cd4*, *Ccr6*) subset was decreased in MLN from RAG2KO mice compared to WT mice. Conversely, clusters 4, 11, and 15 (ILC3/ILC1), as well as cluster 16 (ILC2), were exclusive of RAG2KO mice (**Fig. 2 C-F**). The existence of 4 additional distinct clusters indicates the diversification of ILC subpopulations in RAG2KO mice. ILC3 were comparable in terms of gene expression profiles in both mouse strains (33 DEGs). However, NK cells, ILC1, ILC3/ILC1, and ILC2 were transcriptionally divergent, with 1636, 337, 1277, and 437 DEGs, respectively, between both strains (**Fig. 2 G**). These data highlight the transcriptional divergence in ILCs and NK cells in RAG2KO compared to WT mice.

**Figure 2:**
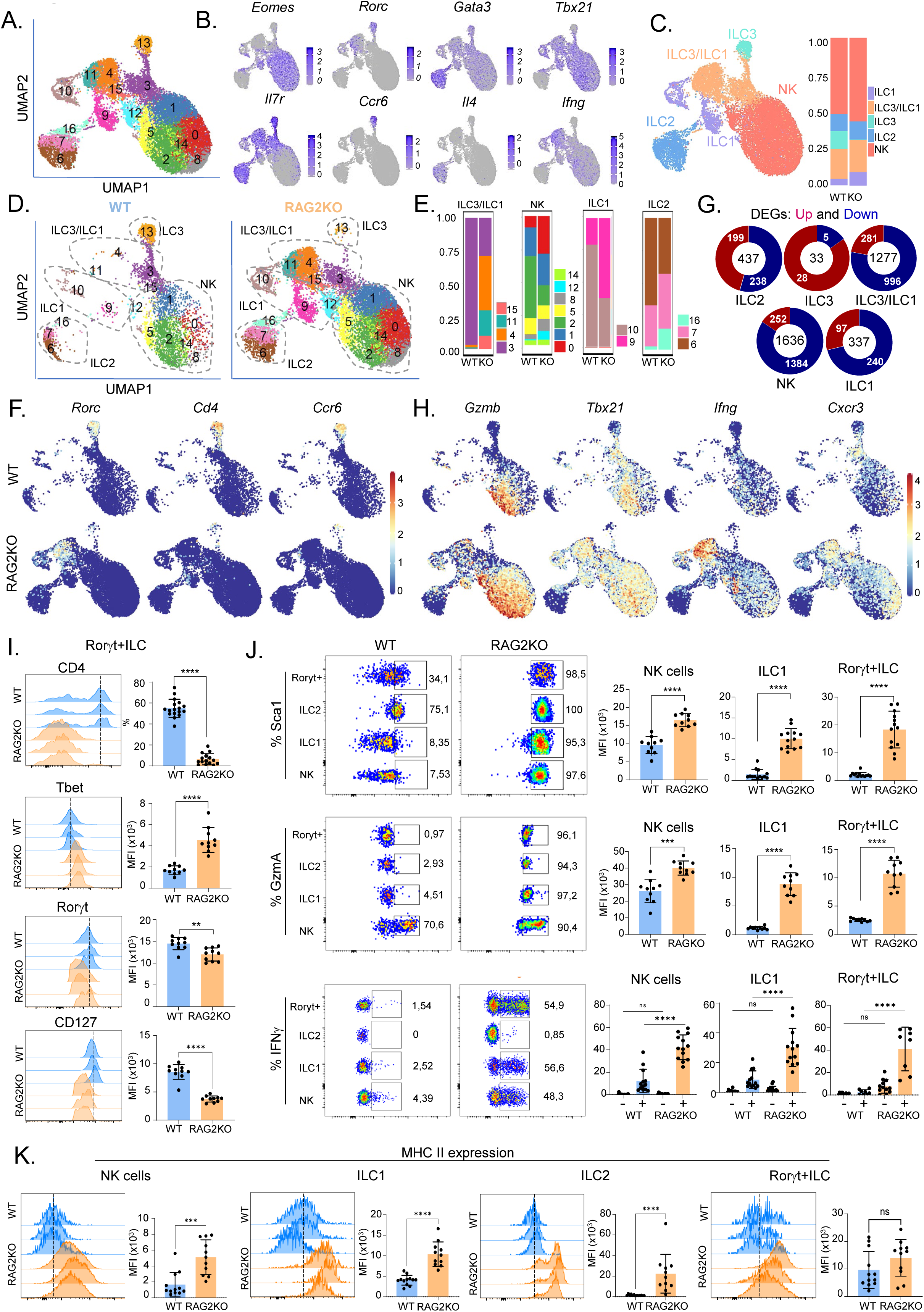
Diversification and distinct functional states of innate lymphocytes in the absence of adaptive IS. scRNA-seq was performed on sorted ILCs and NK cells of MLN from WT and RAG2KO mice. (A) UMAP plot depicting Seurat clusters of sorted innate lymphocytes (ILCs and NK cells) from MLN of WT and RAG2KO mice. (B) UMAP plots showing the expression levels of selected marker genes. (C) UMAP plot displaying identified cell populations and their distribution across WT and RAG2KO genotypes. (D) UMAP plots depicting Seurat clusters split by genotype (WT vs RAG2KO). (E) Cluster representation in each population (ILC1, ILC2, IL3/ILC1 and NK cells). (F) UMAP plot showing the expression of *Rorc*, *Cd4*, and *Ccr6*. (G) Donut plot highlighting the differentially expressed genes (DEGs) between RAG2KO and WT mice, with the total number of DEGs displayed at the center of each plot. (H) UMAP plots showing the expression of specific genes in innate lymphocytes from WT and RAG2KO mice. (I) FACS histograms displaying expression of CD4, Rorγt, Tbet, and CD127 along with MFI of Tbet, Rorγt and Cd127 as well as the percentage of CD4 expression values in Rorγt+ ILCs of MLN. (J) FACS analysis of Sca1, GzmA, and IFNγ expression in ILC3, ILC2, ILC1, and NK cells, presented with MFI values or percentages of expressing cells in each subset. IFN-γ expression is analyzed in two conditions: unstimulated and stimulated with anti-CD40 for 12 hours. (K) FACS histograms showing the expression levels of MHC II on each subset of innate lymphocytes from WT (blue) and RAG2KO (orange) mice, accompanied by MFI. Data (I-K) represent at least two independent experiments and are presented as mean ± SD. Each symbol represents an individual mouse. Data were analyzed using Mann–Whitney tests, except for IFNγ expression in (J), where an one-way ANOVA test was used. Significance is indicated as follows: *P < 0.05, **P < 0.01, ***P < 0.001, ****P < 0.0001; ns indicates not significant (P > 0.05).

In addition, in RAG2KO mice, the expression of cytotoxic molecules (*Gzmb* and *Gzma*) extended beyond NK cells. Furthermore, multiple ILC clusters from RAG2KO mice exhibited expression of genes associated with activation and Th1/ILC1 phenotypes (*Tbx21, Cxcr3,* and *Ifng*) (**Fig. 2 H**). This unrestrained expression of ILC1/Th1 markers indicates a type 1 bias in ILC differentiation in RAG2KO mice. A FACS analysis (**Fig. S2, B**) revealed that Rorγt+ILC barely expressed CD4 in RAG2 KO mice, unlike WT in MLN. In addition, these cells express fewer ILC3 markers such as Rorγt and CD127 (Il7ra) (**Fig. 2 I**). Conversely, these cells acquired the master Th1 transcription factor, T-bet indicating that they may correspond to the ILC3/ILC1 subset within the MLN in RAG2KO mice. In agreement with our scRNAseq data, ILCs and NK cells exhibited an activated phenotype in RAG2KO mice, illustrated by the expression of Sca1, Klrg1, and granzyme A (Gzma), and the production of IFNγ (**Fig. 2 J** and **Fig. S2, C**). Remarkably, even without stimulation, a slight increase in IFNγ production was observed in ILC1 and Rorγt+ILC from RAG2KO mice compared to WT counterparts, indicating a steady-state continuous and basal production of this cytokine. Therefore, these findings reveal that adaptive immunity prevents the diversification and type 1 polarization of innate lymphocytes during homeostasis.

### 4. ILCs and NK cells acquire a marked interferon signature associated with a myeloid feature without adaptive immunity

To elucidate the functional profile of ILCs and NK cells in RAG2KO mice, we conducted a GO analysis of DEGs between WT and RAG2KO ILCs. Numerous pathways were over-represented in NK cells and all ILC subsets in RAG2KO ILCs, including the response to type II IFN, antigen processing and presentation, and cell-killing features (**Fig. S2, D** and **E**). A closer examination of the corresponding transcripts revealed pathways typically associated with myeloid cells, such as response to virus or bacteria (*Gbps*, *Bst2, S100A8),* and antigen processing and presentation (*Psmb8*, *Cd74*, and *H2-Aa)* (**Fig. S2, D-F**). *Plac8*, a transcription factor typically associated with the myeloid lineage, was exclusively enriched in RAG2KO ILCs (**Fig. S2, F**). Accordingly, FACS analysis showed that the monocytic/macrophage markers CD64, CD74, and MHC II were upregulated on the surface of ILCs and NK cells of RAG2KO mice (**Fig. 2 K** and **S2, G-H**). These findings uncover that adaptive immunity prevents ILCs from acquiring some myeloid features and a marked IFNγ imprinting during homeostasis.

### 5. IECs adopt a strong IFNγ signature without adaptive immunity and exhibit a shift in AMP expression

Next, we investigated the impact of the absence of adaptive immunity on colonic IECs. To achieve this, as a first approach, we performed a bulk RNA-seq of FACS-isolated IECs (EPCAM+CD45-DAPI-) from the colon of WT and RAG2KO mice. A Gene Set Enrichment Analysis (GSEA) indicated that inflammatory as well as type 1 and 2 IFN pathways were enriched in RAG2KO IECs, as shown by the elevated expression of *H2-Aa*, *H2-Ab1,* and *Gbps* (**Fig. 3 A** and **B**). Prompted by the disparity in bulk gene expression, we sought to precise whether all subsets of colonic IECs are equally affected. To address this, we conducted scRNA-seq on colonic IECs. Eight cell types were identified based on reported markers: colonocytes, goblet cells, Paneth-like cells, enteroendocrine cells, progenitor cells, transient-amplifying (TA) cells, stem cells, and tuft cells. Using markers such as *Krt19* and *Slc28a2*, we defined early and late colonocyte differentiation stages (Fig. 3 C-E). The colonocyte subset expressing high levels of antimicrobial peptides (AMPs), such as *Lyz1* and *Defas*, were defined as Paneth-like colonocytes (**Fig. 3 D**). The distribution of IEC subsets showed minimal changes between mouse strains, except for the Paneth-like cells that were decreased in RAG2KO mice (**Fig. 3 E**). At the transcriptional level, certain IEC subsets, such as early colonocytes, displayed significant differences between mouse strains (1085 DEGs), whereas other subsets, such as tuft cells or endocrine cells, did not show such divergence (**Fig. 3 F**). A GO analysis indicated an over-representation of IFN pathways and immune signatures (antigen presentation-associated genes *H2-Aa*, *Cd74*, *B2m*, *Psmb8*, and *Psmb9*) as a common feature of RAG2KO IEC subsets (**Fig. 3 G**). Consistent with the upregulation of IFNγ gene expression in RAG2KO mice, the expression of MHC II on the surface of epithelial cells was fourfold times higher compared to WT, as shown by FACS analysis (**Fig. 3 H** and **I**). Therefore, IECs adopt a strong IFNγ signature without adaptive immunity.

**Figure 3:**
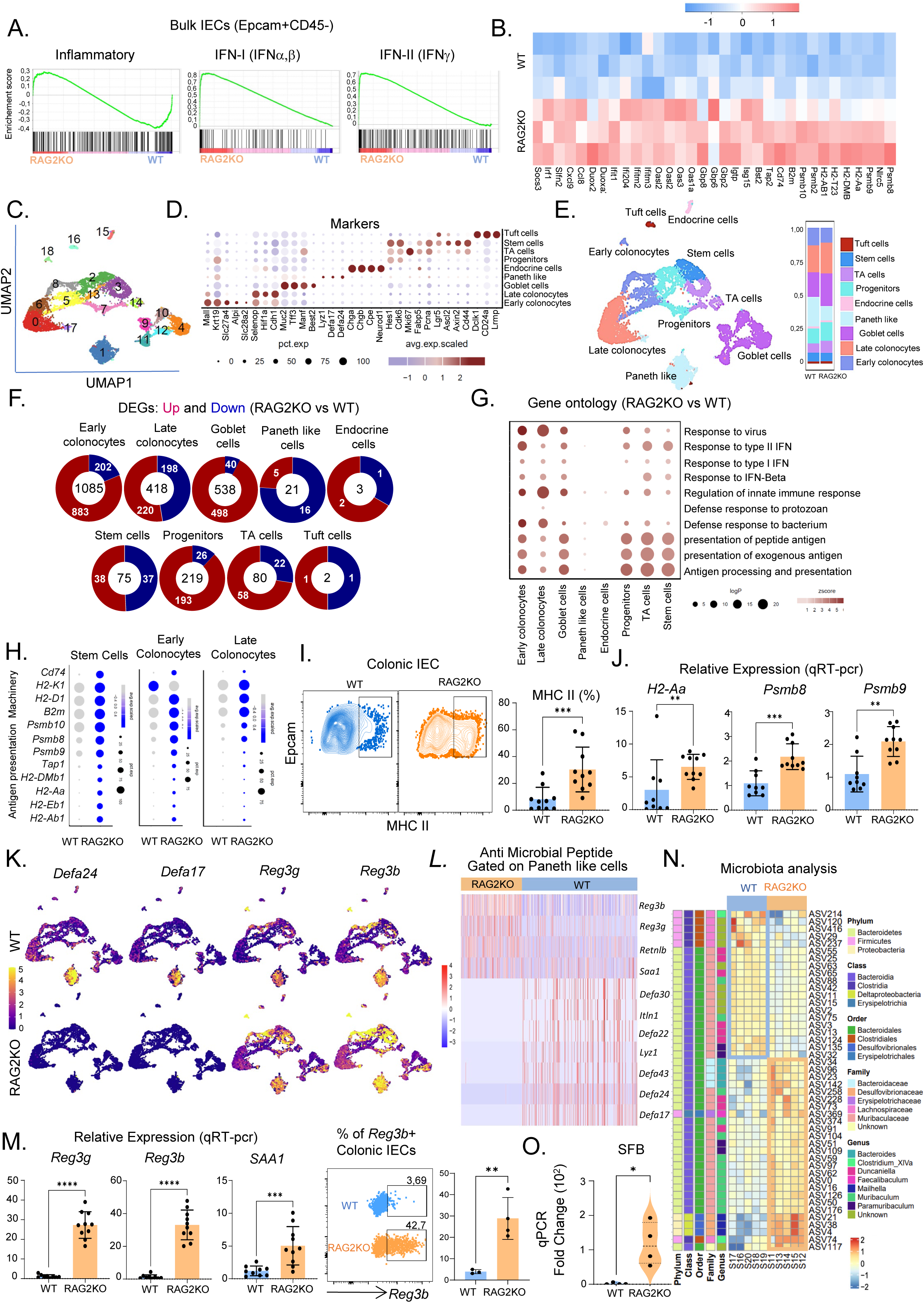
Absence of adaptive IS imposes an IFNγ immune imprinting and an AMP switch on colonic IEC. Bulk and Single Cells RNA sequencing analysis on sorted EPCAM+DAPI-CD45-cells from the colon of WT and RAG2KO mice. (A) GSEA showing enrichment of indicated pathways in bulk transcriptomic data from colonic IEC of RAG2KO (n=3) and WT (n=3) mice. (B) Gene expression heatmap of bulk RNA-seq data for genes linked to IFNγ pathway. (C) UMAP plot of IEC in WT and RAG2KO mice highlighting the identification of 19 distinct clusters. (D) Dot plots illustrating the expression of marker genes used for cluster cell annotation. (E) UMAP showing the name of the identified subsets of IEC and their relative proportions in RAG2KO and WT mice. (F) Donut plot showing the DEGs RAG2KO versus WT with the total number of DEGs displayed at the center of each plot (P value<0.05). (G) Dot plot showing the up-regulated GO terms colored by Z-score. The size of the dot is based on gene count enriched in the pathway (H) Dot plots of indicated gene linked to IFNγ in 3 different cell types, comparing RAG2KO and WT. (I) Representative FACS showing the expression levels of MHC II on colonic IEC cells from RAG2KO mice versus WT associated with the percentage of MHC II positive cells in IEC of WT and RAG2KO. (J) qRT-PCR analysis of the expression of indicated antigen processing and presentation genes in bulk colonic IEC from RAG2KO and WT mice. (K) UMAP showing the expression of indicated AMP genes separated by genotype. (L) Gene expression heatmap showing antimicrobial-related genes in Paneth-like cells of RAG2KO and WT mice. (M) qRT-PCR analysis of the expression of *Reg3y*, *Reg3β* and *Saa1* in bulk colonic IEC from RAG2KO and WT mice alongside representative flow cytometry plot showing *Reg3β* mRNA expression. (N) Heat map depicting differential microbial taxa composition in feces of WT and RAG2KO mice obtained via 16S RNA sequencing analysis. (O) qPCR analysis of SFB abundance in fecal samples from RAG2KO mice compared to WT. Data (I, J, M and O) are presented as mean ± SD. Each symbol represents an individual mouse. Data were analyzed by Mann–Whitney tests (\**P* < 0.05, \*\**P* < 0.01, \*\*\**P* < 0.001, \*\*\*\**P* < 0.0001. ns, not significant (*P* > 0.05).

Notably, RAG2KO mice lacked AMP such as *Defa24* or *Lyz1*, which are typical Paneth markers, while abnormally expressing *Reg3g*, *Reg3b,* and *Saa1*, indicating a marked shift in AMP expression (**Fig. 3 K-M**). Corresponding to this shift of AMP expression, RAG2KO mice exhibited an alteration of the microbiome composition (**Fig. 3 N**). Specifically, RAG2KO mice had more representatives from the Proteobacteria phylum, including pathobionts from the Desulfovibrionaceae and Muribaculaceae families. Conversely, WT mice exhibited a greater representation of Firmicutes species, including Lachnospiraceae (**Fig. 3 N**), suggesting higher production of short-chain fatty acid with a potent immunoregulatory potential (Biddle et al., 2013). We next performed a quantitative PCR (qPCR) analysis of SFB, considering the known influence of these bacteria on the immune system. The results revealed a predominance of SFB in RAG2KO mice (**Fig. 3 O**). In conjunction with the loss of the Paneth-like features, pathobionts and mucin-degrading bacteria emerge in RAG2KO mice. Taken together, these findings uncover that colonic IECs without adaptive immunity adopt a marked IFNγ immune signature and a distinctive AMP expression that likely contributes to the change in microbiota composition.

### 6. Remodeling of innate cell interaction network without adaptive immunity

To explore intercellular communication within the innate cell compartment at homeostasis, we used CellChat to predict the predominant signaling networks (Jin et al., 2021). We identified 57 potential cellular interactions during homeostasis (**Fig. 4 A**). Among these interactions, eight were specific to WT mice, and six were specific to RAG2KO mice. Notably, NOTCH-based interactions in RAG2KO mice were absent (**Fig. 4 A** and **B**). This absence was consistent with the paucity of ILC3 in RAG2KO mice since ILC3s are significant providers of NOTCH ligands such as *Dll1* (Fig. 4 C). In agreement with the regulatory function of Ly6c-MOs (Nahrendorf et al., 2007), we observed that these cells served as the sender and receiver of the immunoregulatory cytokine TGFβ in WT mice. In addition, NK cells constituted a significant source of TGFβ signal for Ly6c-MOs in WT mice. Conversely, this immunoregulatory network was reduced in RAG2KO mice (**Fig. S3, A**). A prominent communication network centered around IFNγ emerged, with ILC3/ILC1 and ILC1 serving as significant producers of IFNγ, while Ly6c+ MOs were the primary recipients in RAG2KO mice. Remarkably, these interactions were not detectable in WT mice under steady-state conditions (**Fig. 4 A** and **B**). The prominent role of IFNγ expressed by ILC3/ILC1 and ILC1 cells in the cellular network in innate cells during homeostasis is further validated by another computational method (**Fig. S3, B**) (Browaeys et al., 2019). The single-cell regulatory network inference and clustering (SCENIC) computational method unbiasedly identified *Irf1* and *Stat1*, transcription factors downstream of IFNγ, as key regulons enriched in Ly6c+ MOs of RAG2KO mice (**Fig. S3, C**) reinforcing the impact of IFNγ at homeostasis without adaptive immunity. Consequently, these data unveil an underappreciated innate homeostatic interaction centered around IFNγ between Ly6c+ MOs and ILC3/ILC1 and a remodeling of the innate cell interactome network during homeostasis in the absence of adaptive immunity.

**Figure 4:**
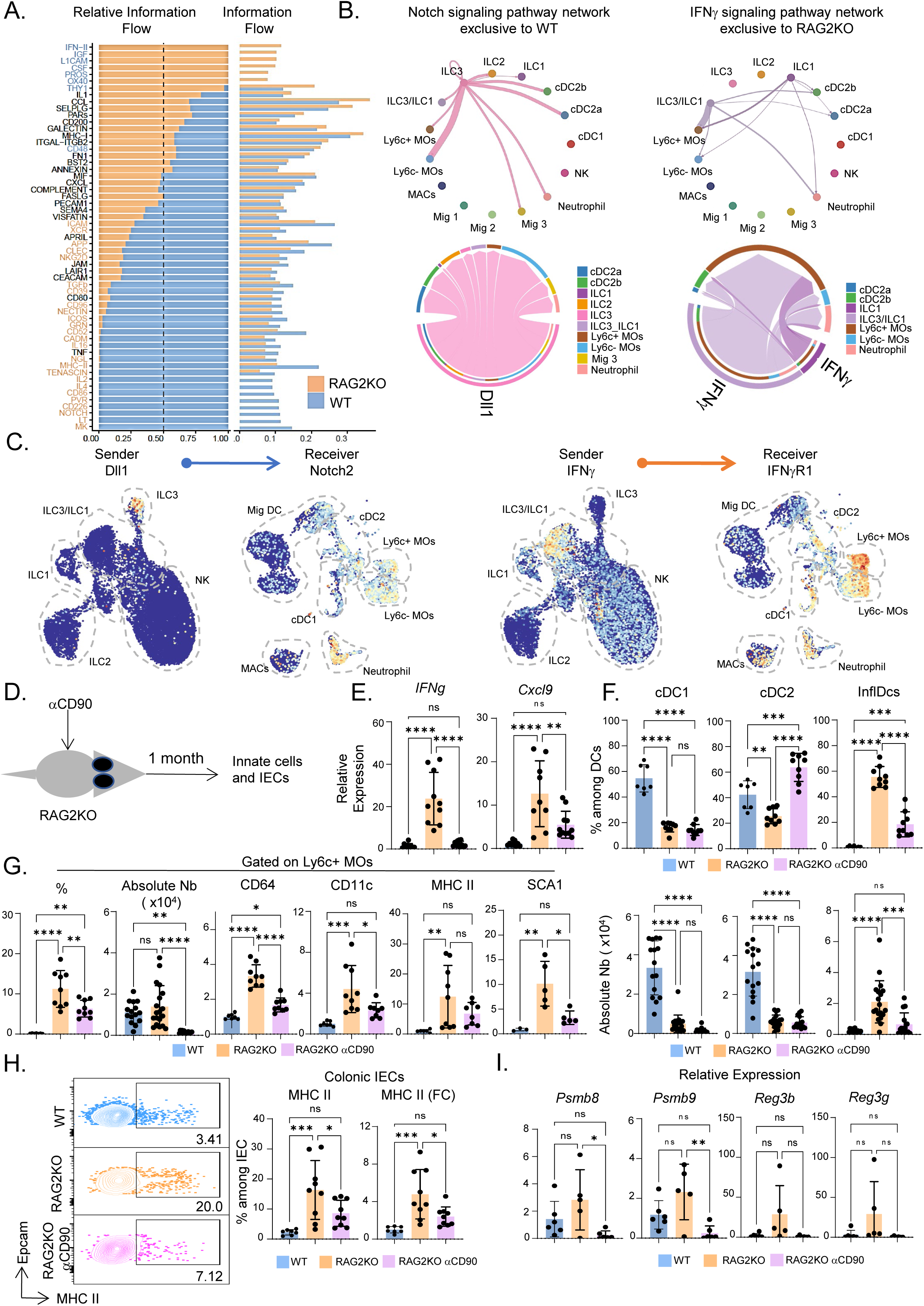
Distinct innate cell interactome network in the absence of adaptive immunity. (A) Information flow charts indicating ranked significant pathways of all innate cells (myeloid and innate lymphocytes) from WT and RAG2KO mice. (B) Circle plots illustrating the inferred NOTCH and IFNγ signaling networks among total innate cells, with edge width representing communication probability. (C) UMAP visualizations depicting the expression of *Dll1*, *Notch2*, *Ifny*, and *Ifnyr1* transcripts in specified innate populations. (D) Shema showing the strategy of innate lymphocyte depletion. (E) Relative expression of *Ifn*y and *Cxcl9* in MLN of WT, RAG2KO, or RAG2KO-depleted mice. (F) Histograms from FACS analysis showing the percentage and absolute numbers of cDC1, cDC2, and InflDCs among total DCs (CD11c+MHC II+) in MLN of WT, RAG2KO, or RAG2KO-depleted mice. (G) Histograms from FACS analysis showing the percentage and absolute numbers of Ly6c+ MOs among CD45+ cells in MLN of WT, RAG2KO, or RAG2KO-depleted mice, with accompanying FACS plots indicating MHC II and CD11c expression. (H) FACS plots of colonic IECs showing MHC II expression, along with histograms of the percentage of MHC II+ IECs and the MFI of MHC II. (I) Relative gene expression (qRT-PCR) of indicated genes in enriched IECs from the colon of WT, RAG2KO, or RAG2KO-depleted mice. Data (E-I) were analyzed using a one-way ANOVA test. Significance is indicated as follows: *P < 0.05, **P < 0.01, ***P < 0.001, ****P < 0.0001; ns indicates not significant (P > 0.05).

To investigate the role of ILCs in this communication network and the functional modulation of innate cells and IECs, we depleted ILCs in RAG2KO mice using anti-CD90 depleting antibodies (**Fig. 4 D**). A reduction in IFNγ and CXCL9 expression was observed in ILC-depleted RAG2KO mice (**Fig. 4 E**). A reduction in InflDCs and Ly6c+ MOs, coupled with a decreased expression of CD64, Sca1, CD11c, and MHC II for the latter, were also observed in ILC-depleted RAG2KO mice (**Fig. 4 F** and **G**), demonstrating the role of ILCs in myeloid differentiation at homeostasis. In colonic IECs, a significant attenuation of AMP (*Reg3b* and *Reg3g*) expression and antigen presentation pathway (*Psmb8* and *Psmb9*) representation, as well as MHC II expression, were observed upon ILC depletion (**Fig. 4 H** and **I**). These data collectively demonstrate that ILCs play a crucial role in producing IFNγ, which directly modulates the myeloid and IEC compartments under steady-state conditions.

To demonstrate the contribution of IFNγ in shaping the phenotype and function of innate and IECs during homeostasis, we generated RAG2KO mice lacking IFNγ receptor (RAG2KOIFNγRKO) and conducted comparative analyses with RAG2KO mice. In RAG2KOIFNγRKO mice, NK cells and ILCs exhibited reduced expression of CD64, MHC II, and Gzma compared to RAG2KO mice, indicating reduced activation and acquisition of myeloid-like features (**Fig.S3 D-F**). Although Ly6c+ MOs were enriched in RAG2KOIFNγRKO mice, these cells expressed lower levels of CD64, MHCII, MHCI, and CD11c compared to RAG2 KO mice (**Fig.S3 G**). Additionally, inflDCs were reduced in RAG2KOIFNγRKO compared to RAG2KO mice (**Fig.S3 H**). This reduction of homeostatic inflDCs shows that IFNγ signaling contributes to their generation in RAG2KO mice. Besides, IECs expressed lower MHC molecules (**Fig.S3 I** and **J**) compared to the RAG2KO. In RAG2KOIFNγRKO mice, the phenotype was intermediate between that of WT and RAG2KO mice for diverse parameters analyzed, indicating a contribution of other IFN types. Nevertheless, these findings collectively indicate that ILC-derived IFNγ contributes to the phenotypic and functional homeostatic shaping of innate immune cells and IECs, and adaptive immunity limits this process at steady-state conditions.

### 7. Dynamic adaptation of innate and epithelial cells during homeostasis

To further demonstrate formally the importance of adaptive immunity in homeostatic shaping innate cell and IEC phenotypes, we restored the adaptive immune compartment in RAG2KO mice by transferring B and T cells (**Fig. 5 A**). To evaluate the successful engraftment of adaptive cells in reconstituted animals, we examined the presence of IgA-coated bacteria in the transplanted mice and assessed potential impacts on the fecal microbiota composition. RAG2KO mice transplanted with adaptive cells exhibited IgA-coated bacteria (**Fig. 5B**). Correspondingly, the quantification of SFB and *Akkermansia muciniphila* in the feces of transplanted animals were decreased, both of which are known to be regulated by the adaptive immune system (Flannigan et al., 2016; Zhang et al., 2015) (**Fig. 5C**). This reduction validated the functional reconstitution of adaptive immunity in the RAG2KO mice. We then assessed the impact on innate immune cells. In the transplanted mice, Rorγt+ ILC exhibited decreased Tbet expression and upregulated Rorγt+ and CD4 markers in MLN (**Fig. 5D**). Additionally, ILCs and NK cells expressed lower levels of Sca1 and GzmA (**Fig. 5 E**). This phenotypic restoration indicates that adaptive immune cells exert control on innate lymphocytes at homeostasis. Similarly, Ly6c+ MOs showed reduced homeostatic accumulation in reconstituted RAG2KO mice, with decreased expression of activated phenotypic markers such as MHC II, CD64, CD11c, and SCA1 (**Fig. 5 F-G**). In addition, the accumulation of InflDCs decreased (Fig. 5 H), and the expression of *Ifny* and *Cxcl9* was reduced in MLN (**Fig. 5 I**). The transfer of adaptive immunity also reduced the expression of MHC II, Psmb8, Psmb9, and RAG-associated AMPs Reg3β and Reg3γ in IEC (**Fig. 5 J-K**). These findings collectively demonstrate that adaptive immunity regulates the functional states of innate cells and IECs during homeostasis.

**Figure 5:**
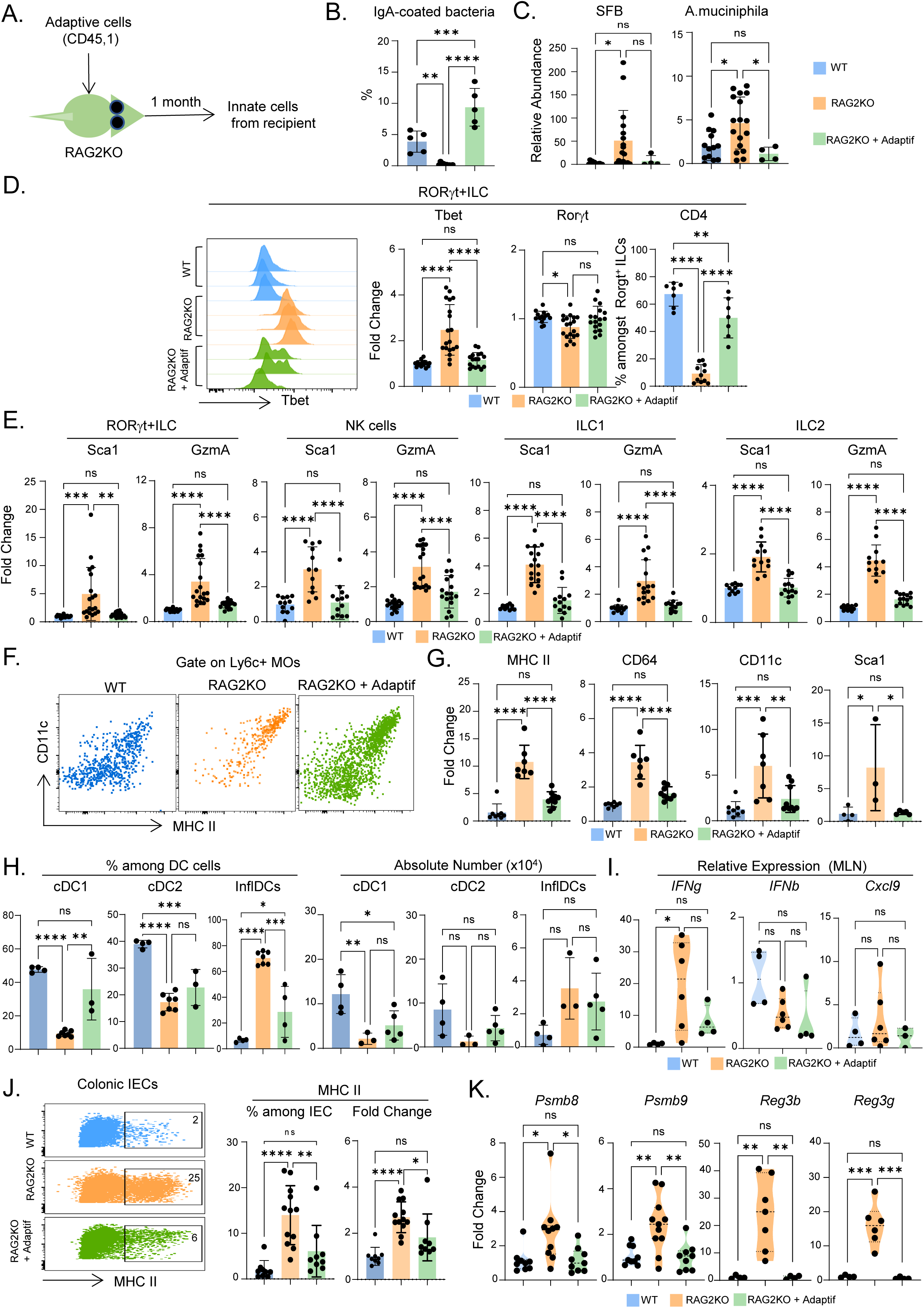
Reversible state of innate immune cells and IECs in presence of adaptive immune cells. (A) Schema showing the adaptive lymphocyte transplantation. (B) Percentage of bacteria coated with IgA in feces of indicated mice. (C) qPCR displaying the relative abundance of SFB and *A. muciniphila* in feces of WT, RAG2KO, or RAG2KO transplanted. (D) Histogram of FACS data showing Tbet expression on Rorγt+ ILCs associated with histograms of the fold change expression of Tbet and Rorγt, and the percentage of cells expressing CD4. (E) Fold change expression from FACS analysis of SCA1 and GzmA across all innate lymphocyte populations. (F) FACS plots showing the expression of MHC II and CD11c on Ly6c+ MOs from the MLN of WT (blue), RAG2KO (orange) and RAG2KO transplanted (green). (G) Fold change expression from FACS analysis of MHC II, CD64, CD11c and SCA1 on Ly6c+ MOs. (H) Percentage among total DC populations and absolute numbers of cDC1, cDC2 and InflDCs in MLN. (I) Relative expression of Ifnγ, Ifnb, and Cxcl9 from qRT-PCR on MLN of WT, RAG2KO and RAG2KO transplanted mice. (J) FACS plot of colonic IEC showing the expression of MHC II associated with the percentage and fold change expression of MHC II expression. (K) Relative expression of indicated genes on enriched IEC from the colon of WT, RAG2KO, or RAG2KO transplanted. Data represent at least 2 independent experiments and are presented as mean ± SD. Each symbol represents an individual mouse. Data were analyzed by Mann–Whitney tests (\**P* < 0.05, \*\**P* < 0.01, \*\*\**P* < 0.001, \*\*\*\**P* < 0.0001. ns, not significant (*P* > 0.05).

## Conclusion remarks and discussion

Innate immune and epithelial cells undergo continuous maturation through intricate microbial and cellular interactions. While these cells are known to shape adaptive immunity (Ivanov et al.; Rie Gaboriau-Routhiau et al.; Brabec et al., 2023), the influence of adaptive immunity on their maturation is less understood. In this study, we reveal that the absence of adaptive immunity largely alters the cellular communication, composition, phenotype and function of innate and epithelial cells during homeostasis. Specifically, without adaptive immunity, innate cells and IECs exhibit a marked type 1 immune signature elicited by ILCs in MLN.

We observed also an increase in the diversity of ILC subpopulations during homeostasis in RAG2KO mice. This diversification involved the emergence of subpopulations/states of ILC2s and NK cells with high levels of MHCII, CD74, and CD64. The latter was initially associated with MO and MAC lineages (Tamoutounour et al., 2013), is also expressed by activated DC2 (Bosteels et al., 2020) and it seems that it can also be expressed by ILCs and NK cells. This is associated with the acquisition of a marker typically linked to myeloid populations, *Plac8*. These specific subpopulations or states of ILC2 and NK cells were rare in MLN of WT mice. Furthermore, we identified a subpopulation of ILC3 with a marked ILC1 phenotype at the expense of the canonical ILC3 subset that expresses a high level of the transcription factor Rorγt. Therefore, our data suggest that adaptive immunity plays a role in maintaining the stability of ILC phenotypes and limiting their diversification under steady-state conditions. The observed diversification of ILC likely represents a dynamic adaptation to the altered local environment to maintain homeostasis. This includes changes in microbial composition, the cytokine milieu, and cellular interactions caused by the absence of adaptive immune cells. Indeed, microbiota and cytokines like IL-12 can drive the transdifferentiation of ILC populations and destabilize canonical ILC phenotypes (Bal et al., 2020; Bernink et al., 2015).

We showed that depleting ILCs reduces the type 1-skewed environment, establishing ILCs as key drivers of this type 1-biased milieu in RAG2KO mice. This type 1 polarization is characterized by overexpression of ISGs, accumulation of a homeostatic inflammatory DCs population, inflDCs with high levels of MHCII, altered AMP expression in IECs, and acquisition of myeloid markers such as MHC II, CD74, and CD64 by ILCs. Blocking IFNγ signaling partially reverses the altered phenotype observed in innate and epithelial cells of RAG2KO mice. This partial rescue indicates that other cytokines, likely other IFNs, also contribute to this type 1 signature bias (Pott and Hornef, 2012; Pott et al., 2011; Saskia Erttmann et al., 2022; Schaupp et al., 2020; Vasquez Ayala et al., 2023). While the roles of type I (IFNα and β) and type III (IFNλ) IFNs in homeostatic shaping are well-established (Pott and Stockinger, 2017; Abt et al., 2012; Van Winkle et al., 2022), our data highlight an overlooked role for type II IFN (IFNγ) in this process during steady-state conditions.

The restoration of a functional adaptive immune cell compartment in RAG2KO mice limits the acquisition of ILC1 markers by ILC3 and reduces this type 1-biased environment in RAG2KO mice. This observation suggests that adaptive immune cells stabilize the ILC3 lineage identity and limit the type 1 skewed milieu. The exact mechanisms by which adaptive immunity can restrict type 1 polarization remain incompletely elucidated. Despite evidence indicating an intrinsic role of RAG gene expression in innate lymphocytes (Karo et al., 2014), our experiments using mixed chimeras, restoration of adaptive immunity, and ILC depletion show that extrinsic factors contribute also to the observed immunological changes in RAG2KO mice. One plausible explanation is that adaptive immunity modulates the composition of the gut microbiota, thereby regulating the expansion of commensal microorganisms that promote type 1 immunity. Accordingly, we have shown that adaptive immunity restoration can reduce the accumulation of potent immunomodulator commensals such as SFB and *Akkermansia muciniphila* (Flannigan et al., 2016; Zhang et al., 2015). However, further research using germ-free and gnotobiotic mice will provide valuable insights into this mechanism. “Homeostatic shaping” is a lengthy process, and several cellular, molecular, and microbial mechanisms likely jointly contribute to it.

Our data underscores that RAG-deficient mice, commonly used to investigate innate cell function or to exclude the implication of adaptive immune cells in diverse settings, exhibit significant alterations in innate and epithelial cells already at steady-state conditions. Innate and epithelial cell divergence at the transcriptome, composition, phenotype, and function levels could bias experimental interpretations. Therefore, our study reminds us that RAG-deficient mice are not only deprived of adaptive immune cells, but their innate and epithelial compartments shift to adapt to the absence of adaptive immunity. Altogether, our findings describe the role of adaptive immunity in influencing the communication, composition, and functional trajectory of innate and epithelial cells during homeostasis and demonstrate their adaptation to maintain homeostasis.

## Material et Methods

### Mice

RAG2KO, WT, and CD45.1 mice were bred and maintained in our animal facilities. To generate RAG2KOIFNγR KO mice, we crossed RAG2KO mice with IFNγRKO mice. All mice were of the C57BL/6J strain and housed in specific pathogen-free (SPF) conditions at the P-PAC facility of the CRCL and the PBES Lyon, France. Unless specified otherwise, both male and female mice, aged between 2 and 8 months, were used for experiments.

### Mix chimera

To generate chimeric mice, 2-month-old recipient RAG2KO mice were irradiated with 600 rads. One 1 week before the irradiation, 100μg of anti-Thy1.2 (clone 30H12 from Bio X Cell) was administered intraperitoneally to eliminate the endogenous NK cells and ILCs. Next, a 1:1 mixture of bone marrow cells isolated from WT B6 and RAG2 knockout mice was injected into the recipients. The chimeric mice were provided with full-spectrum antibiotic water starting 2 days before and continuing for 4 days after irradiation. The mice were analyzed 6 weeks later.

### T and B cell transfer, ILC depletion, and aCD40 injection

RAG2KO mice underwent reconstitution through intraorbital injections of 50 × 10^6 T and B cells derived from the spleen, mesenteric lymph nodes (MLN), peripheral lymph nodes (PLN), and Peyer’s patches of wild-type mice. The cells were purified using a negative selection approach involving biotin-labeled antibodies against Ly6G (Clone 1A8-Ly6g from Invitrogen), Ter119 (clone TER-119 from OZYME), NK1.1(clone PK136 from BD), and CD11b (clone M1/70 from Biolegend), followed by anti-biotin MACS beads (ref 130-090-485). For ILC depletion, RAG2KO mice received intraperitoneal injections of 100μg of anti-mouse Thy1.2 antibody (clone 30H12) once weekly for three consecutive weeks. To induce immune cell activation, both WT and RAG2KO mice received an intraperitoneal injection of 100μg of anti-CD40 antibody (clone FGK4.5/FGK45) or PBS for 15 hours. All antibodies used for in vivo experiments were obtained from Bio X Cell.

### Speen, MLN, and IEC isolation

Spleen and MLN were placed between two layers of nylon mesh (from Sefar) and compressed using the plunger of a 5 ml syringe. Following dissociation, the cell suspension was filtered through a 70-μm cell strainer and centrifuged at 700×g for 5 minutes at 4 °C. After removing the supernatant, red blood cells were lysed using 9 g/L NH4Cl (Sigma Aldrich). The resulting cell pellet was then resuspended in RPMI supplemented with 10% FBS.

After removing the fat, the large intestines were dissected and longitudinally opened, followed by thorough washing in PBS. The intestines were then cut into small pieces and incubated in HBSS containing 5 mM EDTA at 37°C with continuous agitation. After 30 minutes, the cells were separated from the tissue. IEC and intraepithelial lymphocytes were isolated using a 44% Percoll gradient, with IEL settling at the bottom and IEC forming a distinct layer on top.

### Flow cytometry

Cells, prepared as detailed earlier, were first stained with a yellow viability marker, and unspecific binding was reduced using an FcR-blocking reagent (Miltenyi). Then, cells were stained with antibodies (Table below). In vivo, stimulation was conducted for 15 hours with 100μg of anti-CD40 per mouse. Brefeldin A and GolgiStop (BD Biosciences) were added during organ collection and maintained throughout the cell extraction process until the fixation step. After extracellular staining, the cells were fixed and permeabilized using a Cytofix/Cytoperm kit (BD Biosciences) and stained with antibodies targeting IFNγ, Rorgt, Eomes, and Gata3. However, in most cases, following fixation and permeabilization using eBioscience Foxp3/Transcription Factor Staining Buffer Set, cells were stained for these antibodies targeting Gzma, Tbet, Eomes, Gata3, and Rorγt in the case we were not looking at cytokine expression. Flow cytometry data were collected on an LSR Fortessa using DIVA software and analyzed with FlowJo software (all from BD Biosciences).

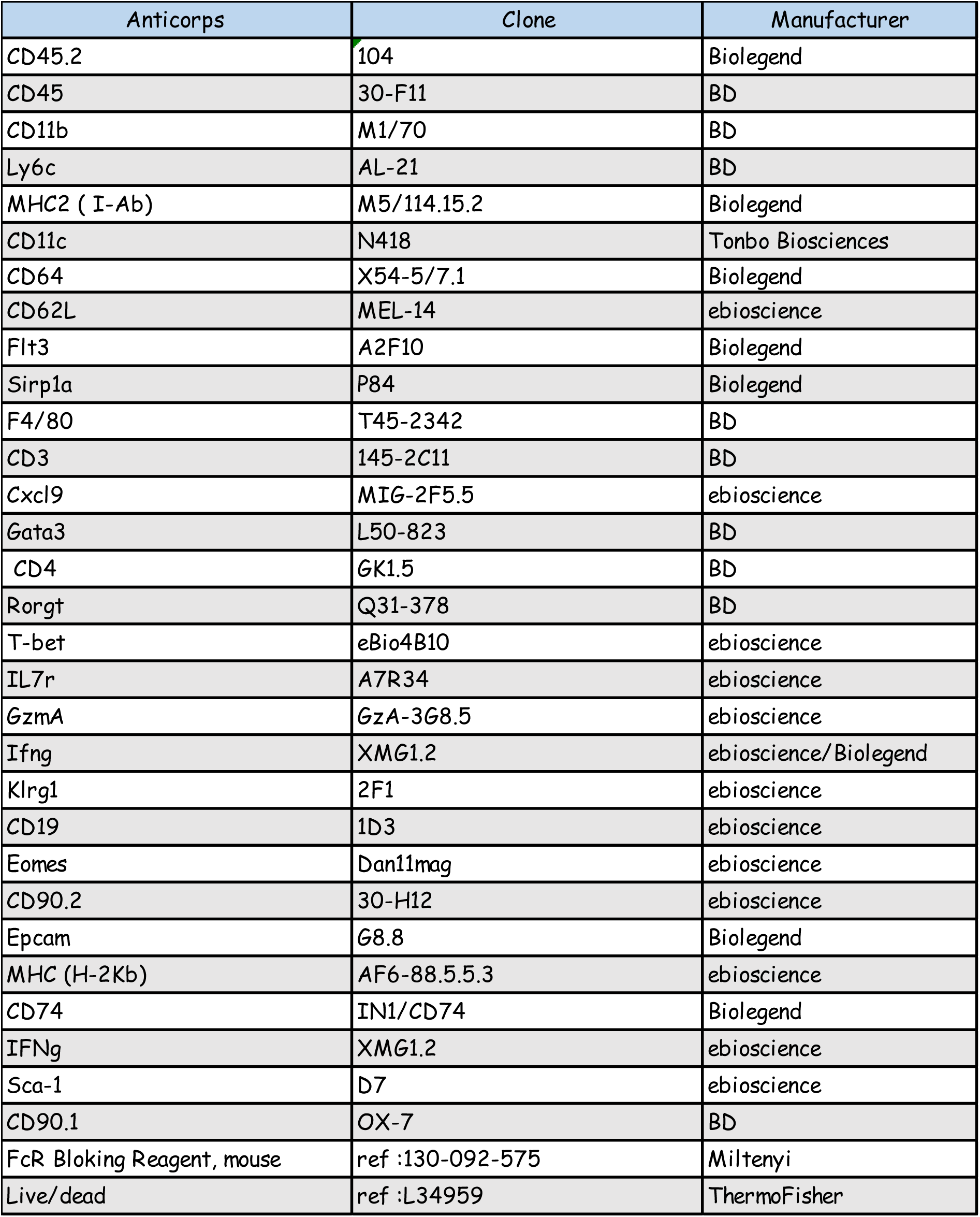

### RT-PCR

RNA extraction was performed using the RNeasy Mini Kit (Qiagen) and reverse transcription with the iScript cDNA Synthesis Kit (Bio-Rad). Quantitative RT-PCR was conducted using LightCycler 480 SYBR Green Master (Roche) and various primer sets on a LightCycler 480 Real-Time PCR System. Samples were normalized to Gapdh and analyzed using the ΔΔCt method. Primer sequences are provided in the accompanying table.

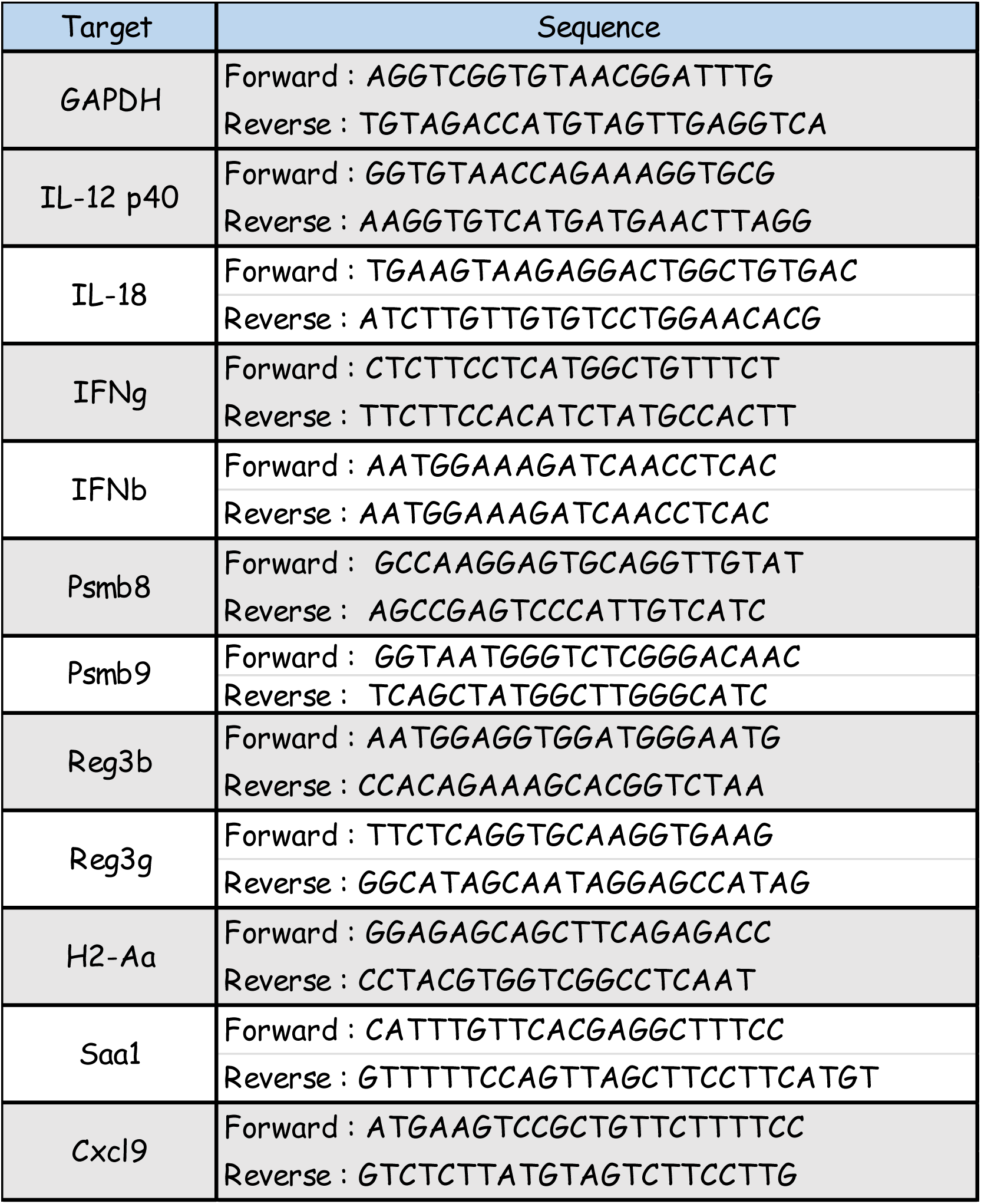

### qPCR and Analysis of fecal microbiota composition through 16S DNA sequencing

qPCR and analysis of fecal microbiota composition through 16S DNA sequencing Bacterial DNA was isolated from mouse fecal samples using the DNA Stool kit (QIAGEN). The 16S rRNA gene was amplified by PCR using primers targeting the V3V4 region (FWD: TACGGRAGGCAGCAG and REV: CTACCNGGGTATCTAAT). PCR conditions and sequencing library preparation were conducted using Metabiote^®^ (Genoscreen, Lille, France). The Amplicon library was sequenced on an Illumina MiSeq using the 2*300 bp V3 kit. Any remaining adapter/primer sequences were trimmed, and reads were checked for quality (≥30) and length (≥250 bp) using cutadapt (Martin, 2011). Reads were further corrected for known sequencing errors using SPAdes (Bankevich et al., 2012) and then merged using PEAR (Zhang et al., 2014). Vsearch pipeline (Rognes et al., 2016) was applied to dereplicate (–derep_ prefix– minuquesize 2) and cluster (–unoise3) the merged reads, as well as check for chimeras (uchime3_denovo). Taxonomic classification of ASVs was performed using the classifier from the RDPTools suite (Trapnell et al., 2014). Sequencing data are deposited in NCBI under the accession number XXXXX. Microbiota composition analysis was done using the R programming language and software (R Development Core Team 2012), specifically using the following packages: ade4 (v1.7-22), vegan (v2.6-4). ASV counts were normalized via simple division to their sample size and then multiplication by the size of the smallest sample. Kruskal-Wallis rank sum tests and post hoc pairwise Wilcoxon rank sum tests were used to detect differences between groups of variables. P values were corrected as necessary using the false discovery rate correction. qPCR reactions were performed with the SYBR Green I qPCR Master mix on the LC480II LightCycler (Both Roche). To ensure specificity, a negative control was implemented involving the use of a water sample instead of the bacterial one. Normalization to Universalis and analysis based on the ΔΔCt method were performed, and primer sequences are provided in the following table.

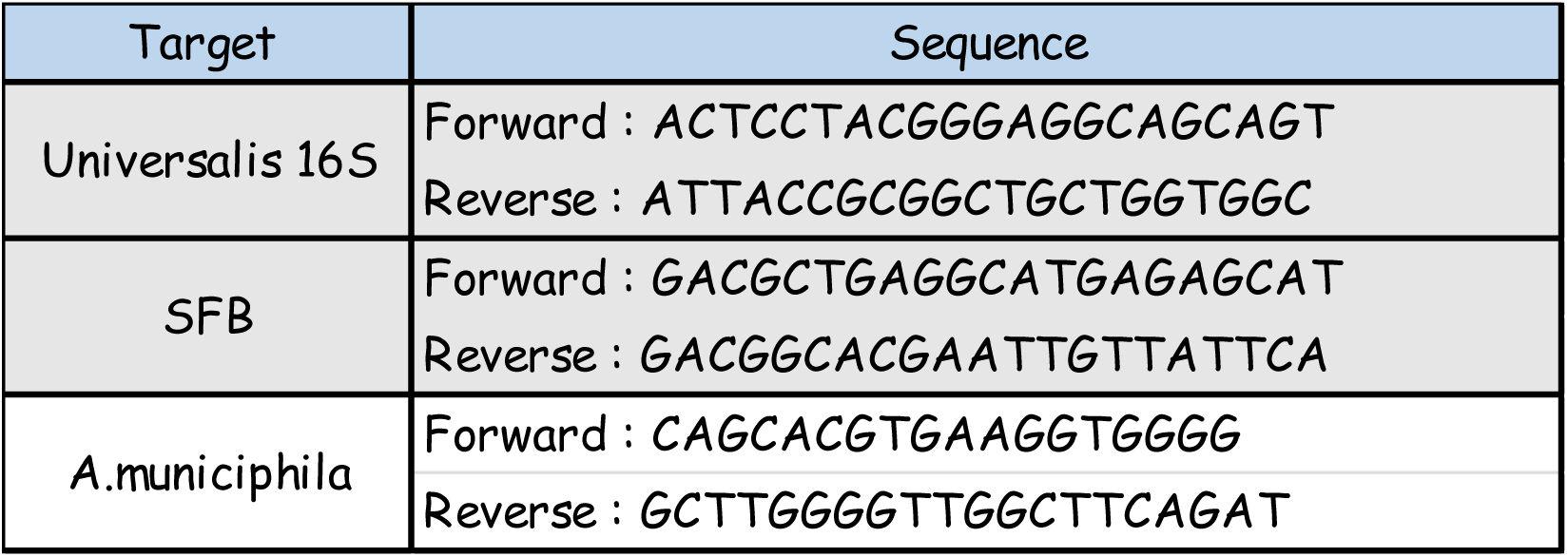

### Elisa

MLN tissues are mechanically disrupted in a RIPA buffer containing a protease inhibitor, utilizing a homogenizer. According to the manufacturer’s instructions, cytokine concentrations in MLN were quantified in pg/mL by bead-based multiplex LEGENDplex™ analysis (LEGENDplex™ Biolegend). Reactions were run in biological duplicate with a BD LSR flow cytometer. The concentrations were processed with LEGENDplex™ Data Analysis Software.

### Bulk RNA-seq

The mesenteric lymph nodes (MLN) and the entire large intestine are homogenized separately in 350 μl and 1 ml of RLT lysis buffer, respectively, using a homogenizer. This is followed by RNA extraction using the Qiagen micro kit. The RNA was then sequenced using Illumina (the Genomic platform of CLB). The data were analyzed using the Galaxy platform (https://usegalaxy.org/). GSEA was performed using the Molecular Signature Database of the Broad Institute (Subramanian et al., 2005).

### Single-cell RNA-seq

Whole colon from WT and RAG2KO mice (littermates, 4-month-old, female, n = 4 per group) were opened by cutting longitudinally through the lumen and washed twice with ice-cold PBS to remove fecal matter end excess fat was removed during the process. The opened colon tissues were washed in a clean petri dish with ice-cold PBS and cut into small pieces. Colonic epithelial cells were enriched following the protocol of isolating mouse intestinal crypts from stem cells to enrich crypt cells (https://www.stemcell.com/how-to-isolate-mouse-intestinal-crypts.html). The cells were sorted by FACS based on EPCAM+DAPI-CD45-with a BD FACSAria™ III sorter (BD Biosciences) instrument. Cells were encapsulated and barcoded based on the standard scRNA-seq protocol (10x Genomics). Single-cell libraries were then sequenced on a NovaSeq System (Illumina).

### scRNA-seq data processing

The Cell Ranger toolkit (version 3.1, 10× Genomics) was used to demultiplex FASTQ files, align to the mouse reference genome (mm10), assign cell barcodes, and generate a unique molecular identifier (UMI) matrix. Seurat (version 4.4.0) (https://satijalab.org/seurat/) was used to analyze gene-cell data matrix ge generated by the cell ranger. Specifically, the raw UMI matrix was processed to filter out genes detected in < 10 cells. The number of genes for each cell was quantified with thresholds of 200-6000 genes to remove empty or doublet cells. Cells that expressed hemoglobin genes 10% were also removed. To screen out cells with high mitochondrial or ribosome proportion, which means low-quality cells, we set the parameters of 10% and 30% to ensure that most of the heterogeneous cell types were included for downstream analysis. We used Harmony (version 1.2.0) (https://github.com/immunogenomics/harmony) to integrate data after normalization to control batch effects. Scrublet (https://github.com/AllonKleinLab/scrublet) and doubletFinder (https://github.com/chris-mcginnis-ucsf/DoubletFinder) were then applied to remove potential doublets with default parameters. We only removed cells that were identified as doublets by both methods, and no cell was filtered out.

### Dimension reduction and unsupervised clustering for scRNA-seq data

Dimension reduction and unsupervised clustering were performed according to the standard workflow in Seurat. To reduce the effect of mitochondrial and ribosome genes on subsequent analysis, a regression parameter (vars.to.regress = c(“percent.mt”,“percent.ribo”) was set in ScaleData() step from the normalized expression matrices. For the clustering, PCA was performed on the top 3000 variable gene matrix generated by the FindVariableFeatures function. The top 15 principal components were used for downstream analyses. For the myeloid and ILC data, we have 34,337 cells in total. After the first round of unsupervised clustering by FindClusters with resolution 0.5, we annotated each cell cluster according to well-established marker gene expression. We identified the major cell types, including myeloid (11,684 cells), ILC (17,232 cells), T (5,299 cells) and B (122 cells) cells. Then, we subset myeloid and ILC cells to do the second-round clustering procedure the same as the first-round clustering. A re-cluster with a resolution of 0.5 was performed for the myeloid cells. 18 clusters were obtained, and clusters 10, 12 and 13 identified as T or ILC cells were excluded from the following analysis. After that, we have 10818 cells in 10 cell types, including Ly6cHigh Monocytes, Ly6c-MOs, MACs, cDC2a, cDC2b, cDC1, Mig1, Mig2, Mig3, and Neutrophiles. From the myeloid Seurat-object, we subset monocytes and cDC2 cells to re-cluster with resolution 0.5 and found two new cell types named MO_DCs and InflDCs. For the ILC cells, a re-cluster with resolution 1.0 was performed. 18 clusters were obtained and no cluster was removed. Gene list scores were calculated using the AddModuleScore function in Seurat. All analyses were performed with Seurat. UMAP visualized clusters with Seurat’s RunUMAP function.

### Differential expression and GO term enrichment analysis

For the identification of differentially expressed genes (DEGs), FindMarkers was performed on all the genes expressed in data slots with a particular parameter logfc.threshold = 0. As recommended, P-value adjustment is performed using bonferroni correction based on the total number of genes in the dataset. The identification of DEGs between two groups of cells was done using the Wilcoxon Rank Sum test (default). P value < 0.05 was considered significant. Function annotation of DEGs was performed by clusterProfiler (version 4.8.3) (Yu et al., 2012). Biological Process (BP) terms were selected, and an adjusted P-value threshold of 0.05 was considered significant.

### Cell-cell communication analysis

Cell-cell communications were estimated by ligand-receptor interactions between cell types using the CellChat (version 1.6.1) (Jin et al., 2021). Due to the need for comparative signal flow analysis between myeloid and ILCs, we combined them and used data and metadata assays to create a CellChat object using the createCellChat function. We analyzed cell-cell communications according to the tutorial on Cellchat. We set the minimum number of cells required in each cell group to 5 in the intercommunication function.

### Pseudotime analyses

Pseudotime analysis and trajectory construction were performed in Monocle2 (version 2.28.0). From the monocytes and cDC2 Seurat-object, we subset Ly6c+ MOs, Ly6c-MO, and MO_DCs cells to be analyzed in pseudotime analyses. Data were converted into a monocle-compatible CellDataSet by newCellDataSet function with default parameters, except lower DetectionLimit = 0.5. Analysis was then performed according to the recommended pipeline. Briefly, the input was created from the UMI count matrix using the newCellDataSet function with default parameters, except expressionFamily = ‘negbinomial.size’. The size factors and dispersion of gene expression were estimated. The tSNE-reduction was reduced based on DDRTree using reduceDimension with special parameters residualModelFormulaStr = “∼nCount_RNA + nFeature_RNA”. Clustering was performed by clusterCells with k=250 to reduce the number of clusters.

### RNA velocity analysis

We used TFvelo (version 1.0) (https://github.com/xiaoyeye/TFvelo) with default parameters for the RNA velocity analysis. TFvelo infers gene dynamics and cell fate based on the abundance of the target gene and its potential transcription factors instead of the relationship between the unspliced and spliced mRNAs. Before velocity analysis, we transferred the Seurat object to the h5ad form to get the input file. The scripts TFvelo_run_demo.py and TFvelo_analysis_demo.py were used to finish this analysis.

### Transcriptional regulon enrichment analysis

We investigated the difference in transcription factors activity using the R version of SCENIC (version 1.3.1). Then, we used the gene count matrix as input for SCENIC. Transcription factors-gene coexpression modules were defined with GENIE3 and refined via RcisTarget using mouse mm9 genome annotation for cis-regulatory analysis. Only genes containing the respective transcription factor binding motif were kept, using gene-motif rankings obtained from https://resources.aertslab.org/cistarget/.

### NicheNet ligand-receptor-target analysis

Using NicheNet R package (’nichenetr’, version 2.0.5, https://github.com/saeyslab/nichenetr) (Browaeys et al., 2019), we cross-validated cell communication results. Ly6c+ MO subpopulations were defined as receivers, and ILCs were defined as senders. In this study, DEG between KO and WT were derived and filtered at a level of adjusted p value (<=0.05) and absolute log2 fold-change (>=0.25). Preconstructed databases were downloaded for ligand_target_martix, ligand_receptor_database, and weighted networks (https://zenodo.org/record/7074291/files/). DEGs in receivers were used to predict the top 30 potential regulatory ligands, target genes on receivers, and the possible cell types as sources of ligand expression.

### Statistics

Statistical analysis was performed using GraphPad Prism. Comparisons among multiple groups were performed using 1-way ANOVA. Data from 2 groups were compared by by Mann–Whitney tests or paired Student’s t-test. P values less than 0.05 were considered statistically significant.

### Study approval

The experiments were conducted following the European Union’s (ARRIVE) guidelines for animal care and validated by the local Animal Ethic Evaluation Committee (AcceS) and the French Ministry of Research.

## Data availability

Data are available in the GEO repository, with accession number:XXXXXXXXXXXXXXXXXXXX.

## Acknowledgments

We thank all the laboratory’s past and present members for their help and discussions, particularly Hélène Taraye, Stéphanie Rodriguez, Manon Pratviel, Maxime Sanchez, and Perla Mteyni. We also thank the CRCL’s platforms, including the flow cytometry, genomic, single-cells and animal facilities (P-PAC and PBES), for advice, help, and assistance. Finally, we thank ANR and ARC for their financial aid.

---

**Figure Supplementary 1:**
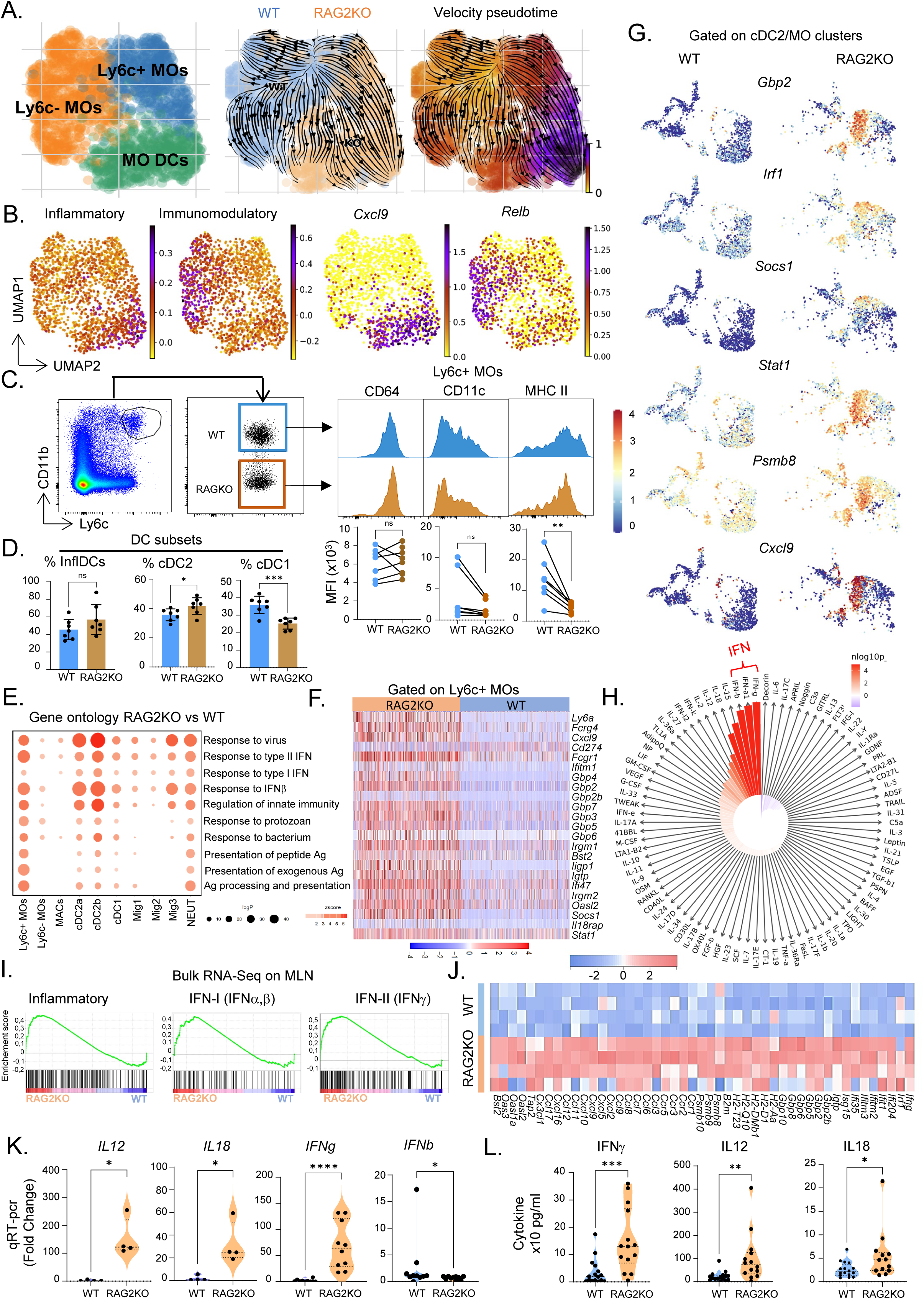
(A) Trajectory inferred by TFvelo: UMAP plots showing (first plot) cell type identification, (second plot) stream plot overlaid on UMAP with WT cells in blue and RAG2KO cells in orange, and (third) pseudotime visualization. (B) UMAP visualization of indicated pathways and genes. (C-D) Mice were irradiated and reconstituted with WT bone marrow (CD45.1) and RAG2KO bone marrow (1:1 ratio). (C) FACS plot showing the gating strategy for Ly6c+ MOs and expression levels of CD64, CD11c, and MHC II (FACS histograms with mean fluorescence intensity, MFI). (D) Distribution of inflammatory DCs (InfDCs), cDC2, and cDC1 cells based on their origin (WT or RAG2KO) in the mixed chimera. (E) Dot plot displaying Gene Ontology (GO) terms for biological processes and molecular functions enriched in RAG2KO mice across all myeloid cells. The dot size represents gene count in the pathway, and the dot color indicates the significance of pathway enrichment. (F) Heat map of some selected IFNγ-induced genes in Ly6c+ MOs. (G) UMAP gated on cDC2 and MO cells showing the expression of selected IFNγ-induced genes. (H) IREA cytokine enrichment plot showing the enrichment score (ES) for each of the 86 cytokine responses in upregulated DEGs of MACs of RAG2KO mice. Bar length represents the ES, shading represents the FDR-adjusted P value (two-sided Wilcoxon rank-sum test), with darker colors representing more significant enrichment (I) GSEA analysis on DEGs RAG2KO versus WT on bulk RNA-seq of MLN. (J) Heat map showing the expression of IFNγ markers on bulk RNA-seq on MLN from WT (n=4) and RAG2KO (n=4) mice. (K) Relative expression of indicated genes from qRT-PCR on MLN. (L) IFNγ, IL12 and IL18 concentration in total MLN. Data represent at least 2 independent experiments and are presented as mean ± SD. Each symbol represents an individual mouse. Data (KL) were analyzed by Mann–Whitney tests and data (C-D) were analyzed by paired Student’s *t* test (\**P* < 0.05, \*\**P* < 0.01, \*\*\**P* < 0.001, \*\*\*\**P* < 0.0001. ns, not significant (*P* > 0.05).

**Figure Supplementary 2:**
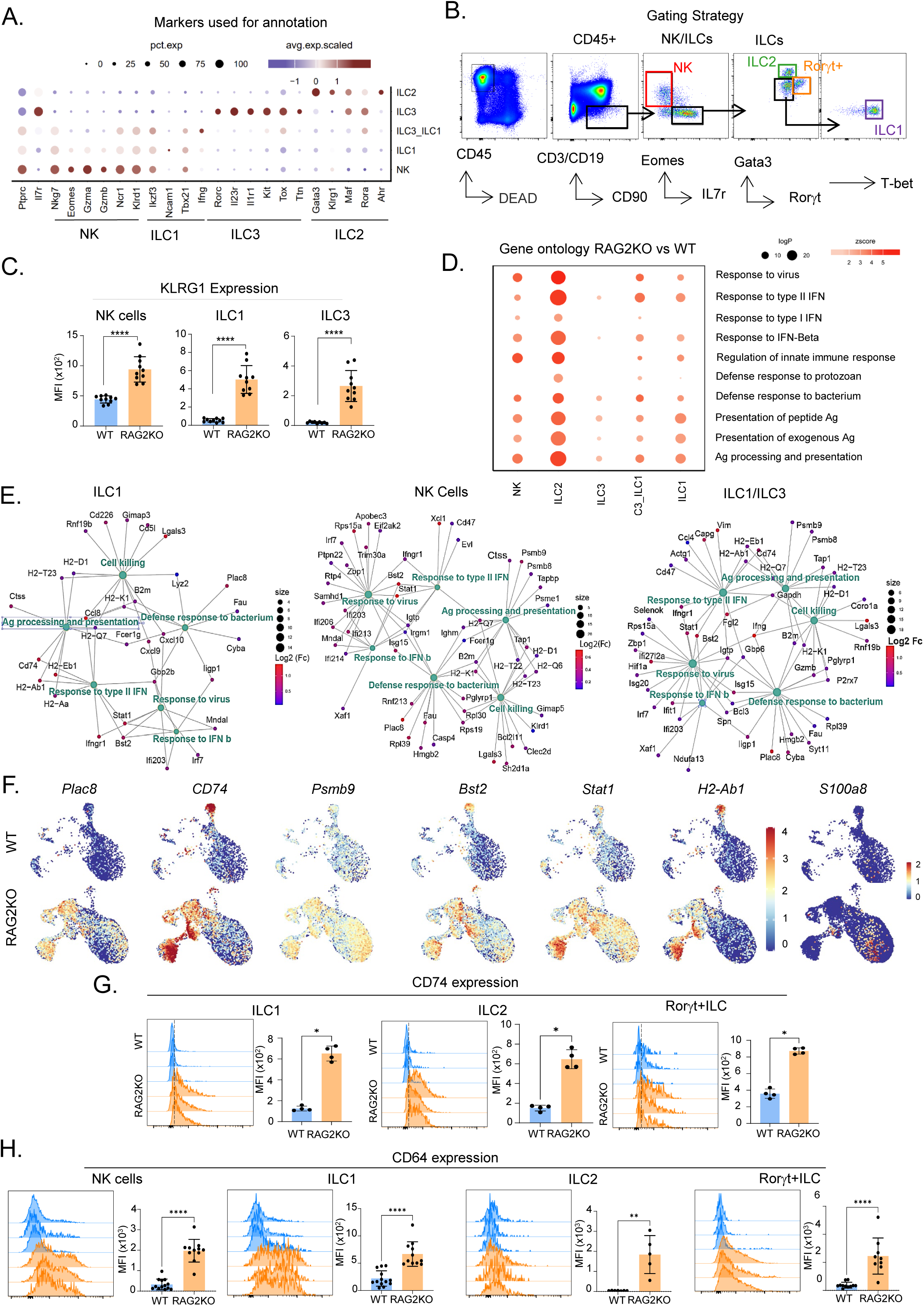
(A) Dot plots showing differential gene expression of selected markers across identified subsets of cells. (B) FACS plots illustrating the gating strategy used for cell identification. (C) MFI of KLRG1 expression on the indicated cell populations. (D) Dot plot representing the enrichment of selected GO terms in upregulated genes within each subset of innate lymphocytes. (E) Cnet plot displaying the association between genes in enriched pathways. The dot color represents the fold change of genes, and the size of the green dots indicates the number of genes enriched in each GO term. (F) UMAP plots depicting the expression of selected genes in WT and RAG2KO mice. (G-H) FACS histograms showing CD74 (G) and CD64 (H) expression on each subset of innate lymphocytes, with associated MFI values. Data represent at least 2 independent experiments and are presented as mean ± SD. Each symbol represents an individual mouse. Data were analyzed by Mann–Whitney tests (\**P* < 0.05, \*\**P* < 0.01, \*\*\**P* < 0.001, \*\*\*\**P* < 0.0001. ns, not significant (*P* > 0.05).

**Figure Supplementary 3:**
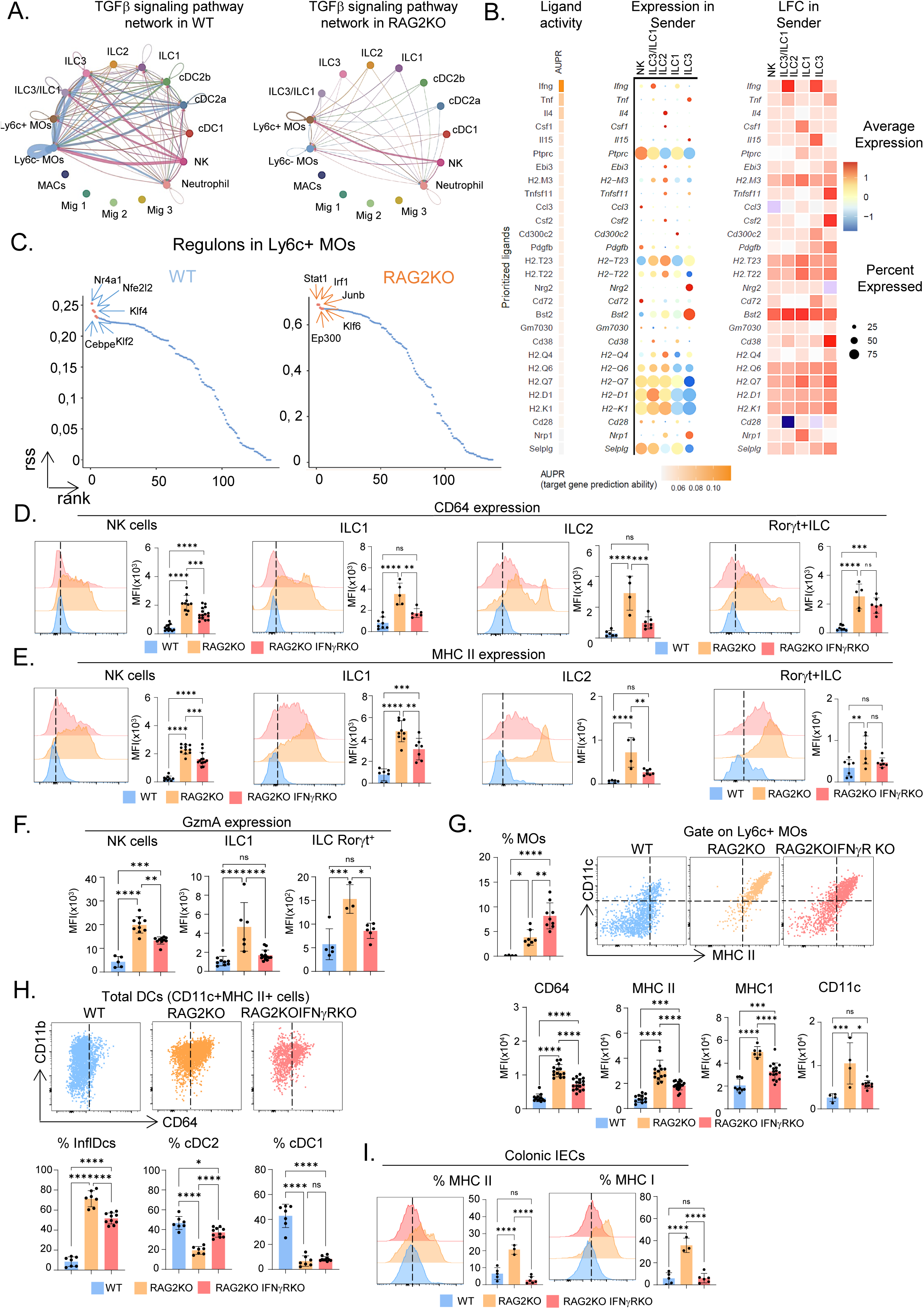
(A) Circle plot illustrating the cell-cell communication network, showing the inferred TGFβ signaling interactions among all innate cell populations. (B) NicheNet analysis showing the ligand-receptor matrix. (C) Scatter plot ranking regulons in Ly6c+ MOs from WT and RAG2KO mice, based on the regulon specificity score (RSS) determined by SCENIC. The top 5 regulons are highlighted. (D-E) Histograms of FACS data showing MHC II and CD64 expression on ILCs and NK cells, along with MFI values from WT, RAG2KO, and RAG2KO IFNγRKO mice. (F) Histograms displaying the percentage of cells expressing Gzma within NK cells, ILC1, and Rorgt+ cells. (G) Histogram showing the percentage of Ly6c+ MOs among all CD45+ cells in the MLN, along with FACS plots of CD11c and MHC II expression on Ly6c+ MOs. Below are histograms showing the MFI of CD64, MHC II, MHC I, and CD11c on Ly6c+ MOs. (H) FACS plots showing CD11b and CD64 expression on total DC cells in the MLN of WT (blue), RAG2KO (orange), and RAG2KO IFNγRKO (red) mice, along with the percentage of cDC1, cDC2, and inflammatory DCs (InflDCs). (I) FACS plot showing the expression of MHC II and MHC I, along with the percentage of MHC II and MHC I bright cells in colonic intestinal epithelial cells (IEC). Data (D-J) represent at least 2 independent experiments and are presented as mean ± SD. Each symbol represents an individual mouse. Data were analyzed by Mann–Whitney tests (\**P* < 0.05, \*\**P* < 0.01, \*\*\**P* < 0.001, \*\*\*\**P* < 0.0001. ns, not significant (*P* > 0.05).

